# Sequence dependencies and mutation rates of localized mutational processes in cancer

**DOI:** 10.1101/2021.10.27.465848

**Authors:** Gustav Alexander Poulsgaard, Simon Grund Sørensen, Randi Istrup Juul, Morten Muhlig Nielsen, Jakob Skou Pedersen

**Affiliations:** Department of Clinical Medicine, Aarhus University, Palle Juul-Jensens Boulevard 82, 8200, Aarhus N, Denmark; Department of Molecular Medicine (MOMA), Aarhus University Hospital, Palle Juul-Jensens Boulevard 99, 8200, Aarhus N, Denmark; Bioinformatics Research Center (BiRC), Aarhus University, 8000, Aarhus C, Denmark

**Keywords:** pan-cancer, mutational processes, hotspots, mutation rate

## Abstract

**Background:** Cancer mutations accumulate through replication errors and DNA damage coupled with incomplete repair. Individual mutational processes often show strong sequence and regional preferences. As a result, some sequence contexts mutate at much higher rates than others. Mutational hotspots, with recurrent mutations across cancer samples, represent genomic positions with elevated mutation rates, often caused by highly localized mutational processes.

**Results:** We analyze the mutation rates of all 11-mer genomic sequence contexts using the PCAWG set of 2,583 pan-cancer whole genomes. We further associate individual mutations and contexts to mutational signatures and estimate their relative mutation rates. We show that hotspots generally identify highly mutable sequence contexts. Using these, we show that some mutational signatures are enriched in hotspot sequence contexts, corresponding to well-defined sequence preferences for the underlying localized mutational processes. This includes signature 17b (of unknown etiology) and signatures 62 (POLE), 7a (UV), and 72 (linked to lymphomas). In some cases, the mutation rate increases further when focusing on certain genomic regions, such as signature 62 in poised promoters, where the mutation is increased several thousand folds over the overall data set average.

**Conclusion:** We summarize our findings in a catalog of localized mutational processes, their sequence preferences, and their estimated mutation rates.

## Introduction

Mutational signatures representing mutational processes have been identified and cataloged through analysis of large cancer genomic data sets. Some mutational processes show strong preferences for certain sequence or regional contexts, not captured by traditional mutational signature analysis. They cause variation in the mutation rate along cancer genomes with some positions displaying dramatically elevated mutation rates. These positions may manifest as mutational hotspots, which are recurrently mutated across cancer patients. Here, we use mutational hotspots identified across 2,583 whole cancer genomes to discover and characterize localized mutational processes, including their mutation rate and sequence dependency.

Cancer arises through an evolutionary process within the body, where cells accumulate somatic mutations throughout life [1, 2]. Consequently, the cancer genome represents a record of the mutational processes that have shaped it since the formation of the zygote. While the majority of mutations are neutral passengers, which do not impact the cellular phenotype, some driver mutations are under recurrent positive selection across many patients and may lead to mutational hotspots [3, 4]. However, the far majority of driver hotspots reside in the protein-coding regions [5]. Therefore, we focus on non-coding regions in the PCAWG dataset [6], where few drivers are expected [7] and where we hypothesize most hotspots are explained by localized mutational processes.

Mutagenesis is a multi-step process starting with either replication error or DNA damage, imperfect DNA repair, and then manifests through replication as mutations in descendent cells [8, 9]. Lesions are frequently formed from endogenous processes, such as the spontaneous deamination of cytosine to uracil, and the majority are successfully repaired by the DNA damage response (DDR) system [10]. Similarly, for lesions from exogenous mutagens, such as those found in tobacco smoke, the vast majority is cleared [11, 12]. Excessive lesion formation may overwhelm the DDR system and result in an increased mutation rate [13, 14].

Mutational processes act with varying intensities across the genome [11, 15–23] and certain sequence motifs experience dramatically elevated mutation rates. This is for instance the case for mutations induced by UV radiation (UV), which preferentially fall in TTT<u>C</u>ST (S=C|G) contexts as C>T mutations [21, 24–30], and certain members of the Apolipoprotein B mRNA Editing Catalytic Polypeptide-like (APOBEC) family of DNA-editing enzymes, which induce high loads of C>T and C>G mutations in T<u>C</u>W (W = A | T) contexts [18, 31–38]. In addition, the APOBECs specifically target single-stranded regions of DNA-level stem-loop structures to produce strand-coordinated clusters of localized hypermutation, as discovered from highly context-specific mutational hotspots [36, 38–40]. Likewise, we may study other localized mutation processes through systematic analysis of hypermutable sites and their contexts across cancer genomes.

Recent large whole-genome sequencing (WGS) datasets have powered landmark discoveries of mutational processes [6, 11, 41, 42]. Mutational signature analysis has been a key tool for disentangling the mutational processes shaping these genomes [11, 18, 22, 43]. It exploits that mutational processes are shared across patients, though with varying intensities. Using non-negative matrix factorization (NMF), recurring profiles of mutation types and contexts that represent individual mutational processes are identified and their exposure in each genome evaluated [11, 18, 43].

Given the high number of free parameters and limited data availability, mutational signature analysis was only recently expanded from considering trinucleotide (±1 base pair [bp] neighbors) contexts to pentanucleotide contexts (±2 bp) [22, 44]. Some mutational processes may further depend on regional properties such as chromatin-organization [45–47], transcriptional activity [11, 48–50], and replication asymmetry [51, 52]. As all mutations are weighted equally, traditional signature analysis has limited power to learn the extended sequence contexts and regional preferences of rare localized mutational processes, which are generally underexplored [53].

We here aim to characterize the sequence-dependency and mutation rate of localized mutational processes. We categorized all single base substitutions based on their extended sequence contexts, by considering their five bp up- and down-stream regions (11-mers). This allowed us to evaluate the mutation rate for different sequence contexts. We then associated context-based categories of mutations with mutational signatures and their associated mutational processes. By exploiting that hotspots pinpoint sequence contexts with elevated mutation rates, we identified localized mutational processes and characterized their sequence and genomic feature preferences. Based on this, we decompose the factors that increase the mutation rate in increasingly smaller parts of the genome and evaluate how these factors explain the elevation in mutation rate. We contribute a comprehensive pan-cancer catalog of localized mutational processes associated with mutational signatures.

## Results

### Baseline mutation rate across families of 11-mers

To estimate mutation rates, we initially identified 343,923 coding and 41,318,716 non-coding single nucleotide variants (SNVs) from the PCAWG set of 2,583 whole cancer genomes [6] (**Fig. 1a)**. Our analyses focused on the non-coding SNVs, which occur at an overall mutation rate of 5.96 SNV/patient/Mb (baseline mutation rate) across the dataset.

**Figure 1.**
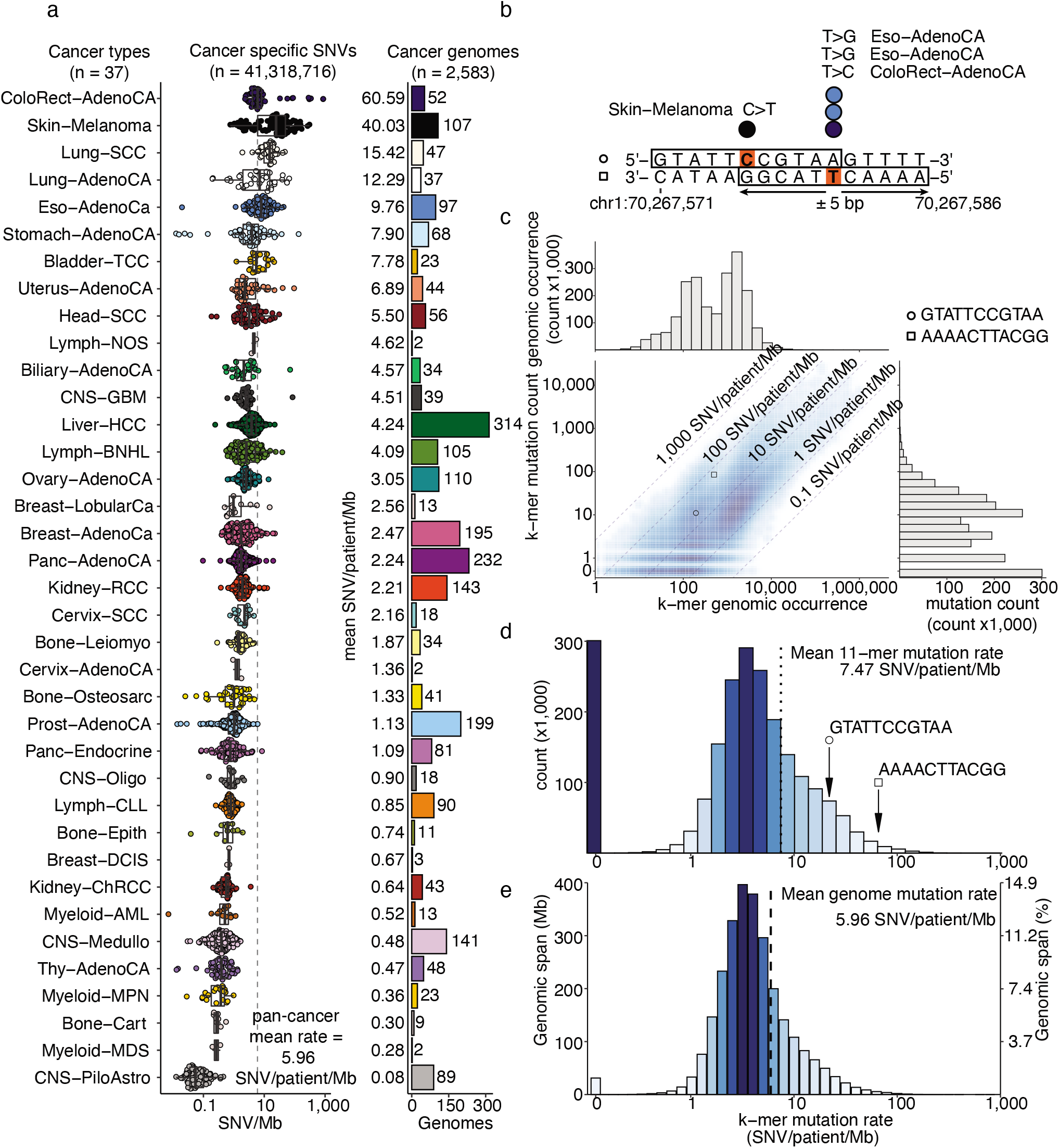
Mutation data and differential mutability of 11-mers. (a) Non-coding mutation rate (left) and the number of cancer genomes (right) grouped and colored by cancer type. Fig. 1a may be regarded as the color legend for cancer types throughout all figures, (b) Illustration of singleton and hotspot single nucleotide variants (SNVs). Strand symmetry is assumed in the analysis and mutated base pairs are represented by their reference pyrimidines (orange). Mutations are annotated with the ±5 bp nucleotide context on the strand of the mutated pyrimidine and represented as 11-mers (framed) in the downstream analysis, (c) Distributions of 11-mer (n=2,097,090) reference (hg19) occurrence (x-axis) and pan-cancer mutation count (y-axis). (d) K-mer count distribution of 11-mer mutation rates, (e) Genomic occurrence (span) distribution of 11-mer mutation rates. K-mer span = 2,684,570,106 bp (100%)

To investigate the sequence dependency of mutations, we classified all genomic positions (n=2,684,570,106) by their 5 bp up- and downstream context, which we considered as 11-mer sequences (**Fig. 1b**). To achieve strand symmetry, base pairs were viewed from the strand that contains the pyrimidine. Hence, 11-mer sequences representing genomic positions with a purine on the plus strand were reverse complemented.

The human genome (hg19) contains 2,097,090 unique strand-symmetric 11-mer sequences. Each 11-mer represents a family of concrete instances along the genome, with some families much larger than others. Unless otherwise stated, we will refer to 11-mer families simply as 11-mers. For each family, we calculated the average mutation rate per patient across the dataset, for example the AAAAC<u>T</u>TACGG family has a mutation rate of 65.8 SNV/patient/Mb, and constitutes 500 instances with 85 SNVs across the 2,583 patients (**Fig. 1c**).

We chose to base our analysis on k-mers of length 11 as they provided an extended mutational context while allowing for a sufficient number of expected mutations for each family of k-mer (19.7 SNVs per 11-mer) to achieve useful mutation rate estimates (Methods; **Suppl. Table 1**).

### Highly variable 11-mer mutation rate

We observed a mean mutation rate of 7.47 SNV/patient/Mb across all families of 11-mers, with a high degree of variation (sd 13.1). 14.3% of 11-mers (*n* = 300,837) harbor no mutations at all, while the rest (85.7%; *n* = 1,796,253) have mutation rates ranging from 0.12 to 774 SNV/patient/Mb, displaying a 6,492-fold difference. This high variation illustrates the inherent heterogeneity of the mutation rate of 11-mers across the genome **(Fig. 1d)**.

When we weigh 11-mer mutation rates by their number of genomic instances, we recover the baseline mutation rate (5.96 SNV/patient/Mb; **Fig. 1e**). In the downstream analyses, we focus on these weighted mutation rates to allow comparison between different genomic subsets.

Some of the variation in mutation rates is a consequence of the sampling variation caused by differences in 11-mer family sizes (i.e. their genomic spans) (**Fig. 1e**). Given uniform sizes, each family would span 1,280 instances. However, the observed number of instances per family range from 1 to 4,674,610 (median 608). For instance, non-mutated 11-mers (14.3% of all) only span 1.2% (31.2 Mb) of the genome, as most are represented by a small number of instances (median 83). Similarly, there are 1.5% highly mutated 11-mers (≥50 SNV/patient/Mb; *n* = 32,080), which only span 0.6% (16.7 Mb) of the genome and thus also represent smaller than average 11-mer families (median 128), though to a less extreme degree. However, the variation in family size is much greater for the highly mutated group than the non-mutated group (sd 2,075.4 vs 101.4).

Although many of the highly mutated 11-mers are rare, some of them are not. Common 11-mers, with equal to or more instances than the median (≥608), make up 9.4% (*n* = 3,023) of all the highly mutated 11-mers (*n* = 32,080). Thus, the high degree of variation in mutation rates across 11-mers does not appear to be governed by family size alone. Consistent with prior findings [30, 54–56], we expect that some of the variability is explained by highly mutable extended contexts.

### Assignment of mutated 11-mers to mutational processes

We next sought to identify and group mutated 11-mers by their underlying mutational processes, to characterize their relative mutation rates and extended sequence preferences. As a proxy for mutational processes, we used the 60 mutational signatures from the PCAWG consortium, generated using the SignatureAnalyzer software [11, 18, 22, 43].

Cancer genomes were grouped into cohorts with shared signature exposure (≥5% exposure; Methods), allowing us to study 11-mers across genomes with potential for shared mutational processes. We obtained 57 signature-exposed cohorts (**Fig. 2a**) each representing between 1 and 2,049 genomes inferred to share a mutational process either pan-cancer or cancer type-specific (**Fig. 2b**). As the mutation burden of a cancer genome is typically explained by multiple signatures, the signature-exposed cohorts also overlap in their ascribed genomes. Consequently, some genomes are members of several signature-exposed cohorts.

**Figure 2.**
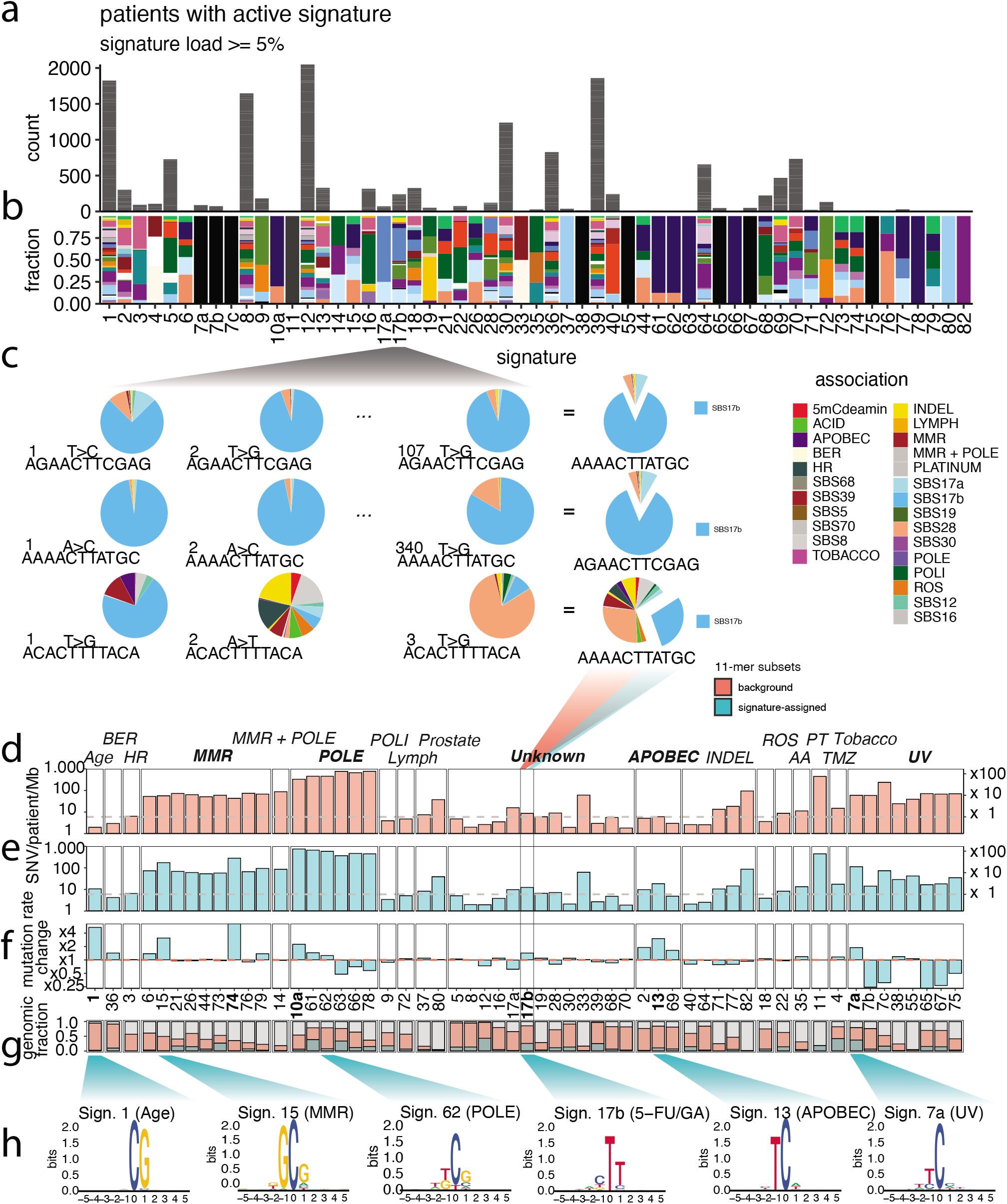
Assignment of cohorts and 11-mers to mutational signatures. (a) Stratification of genomes based on mutational signature load into 60 so-called activity cohorts. Each activity cohort comprises a number from 0 to 2049 genomes (median 48). Cancer type color legend can be found in Figure 1a. (b) Fraction of cancer types in each activity cohort, (c) 11-mer counting across the genomes subjected to signature X. Each mutated position has a probability distribution of possible explanatory signatures. The average signature probabilities across identical 11-mers make up a probability distribution for each unique 11-mer. Hard assignment of an 11-mer to a signature is based on the max probability in the signature distribution, (d) Mean mutation rate of (red) all mutated 11-mers in an activity-cohort. The mean mutation rates (left y-axis) are compared to the global mutation rate (grey dashed line) and represented with a fold-change (right y-axis). Some of the signature-stratified subsets of 11-mers have mutation rates far exceeding the global rate, thus these signatures are extremely active in a subset of contexts, (e) Mean mutation rate of (blue) signature-assigned subset of 11-mers. The mean mutation rates (left y-axis) are compared to the global mutation rate (grey dashed line) and represented with a fold-change (right y-axis). (f) Fold-change in mutation rate from (red dashed line) all mutated 11-mers in an activity-cohort to (blue) signature-assigned subset of 11-mers, (g) Fraction of the genome spanned by 11-mers selected in each analysis step, (h) Sequence logos show emerging context from the subset of signature-assigned 11-mers, mostly capturing contexts evident from the signatures’ profiles.

Some processes were exclusive to distinct tissues, such as UV exposure to the skin (signature 7a; 89 melanoma genomes), while other widely active processes of unknown etiologies, such as signature 17b, possibly related to gastrointestinal cancer or 5-fluorouracil exposure, were found across many cancer types (240 genomes, 13 cancer types). The intrinsic clock-like process of 5-methylcytosine deamination (signature 1) was active in the far majority (70.7%) of all genomes (1,825 genomes, 37 cancer types).

From the 11-mers in each signature-exposed cohort (**Fig. 2c**), we computed the cohort-wise mutation rates (**Fig. 2d**). As expected, we observed that some of these signature-exposed cohorts had much elevated mutation rates compared to the pan-cancer baseline mutation rate, including cohorts defined by signatures associated with mismatch repair (MMR; 63.5 ± 13.2 SNV/patient/Mb; 10.7x), POLE (579.5 ± 183.9; 97.4x), and UV (79.2 ± 66.6; 13.3x) (**Fig. 2d**; **Suppl. Fig. 4**).

For each signature-exposed cohort, we next identified the subset of 11-mers that can be explained primarily by the defining signature. We use the probabilities that individual signatures generated the observed mutations to assign 11-mers to their explanatory mutational process (**Fig. 2c**; Methods).

We characterized the mutation rates of these signature-assigned 11-mers, and found that the rates of a number of signatures were much higher than both the baseline (**Fig. 2e; Suppl. Fig. 4**) and previous analysis step (**Fig. 2f**), most notably signatures related to UV (7a), APOBEC (13), MMR deficiency (74), and POLE deficiency (10a). The 11-mers ascribed to signatures of age, MMR, POLE, and APOBEC generally spanned low fractions of the genome (2-8%). While the 11-mers assigned to tobacco, UV, and signature 17b, spanned large fractions of the genome (42%, 37%, and 26%, respectively; **Fig. 2g**).

We evaluated sequence preferences as logo plots relative to the genomic base composition (**Fig. 2h**) and relative to the composition dictated by the mutational signature (Methods; **Suppl. Fig. 4, Suppl. Fig. 5**). We observed that the base composition in the signature-assigned 11-mer sets mostly recapitulated the composition expected from the signature.

### Hotspots identify 11-mers with high mutation rates

We consider hotspots as proxies for highly mutable positions in the genome. We hypothesize they may be targeted by highly localized and hence context specific mutational processes, which we aim to characterize. From recurrently mutated positions (**Fig. 3a**), we identified 2,842,934 SNVs across 1,339,497 hotspots in the non-coding part of the genome and 17,856 SNVs across 8,173 hotspots in protein-coding regions (**Fig. 3b**) [5, 7].

**Figure 3.**
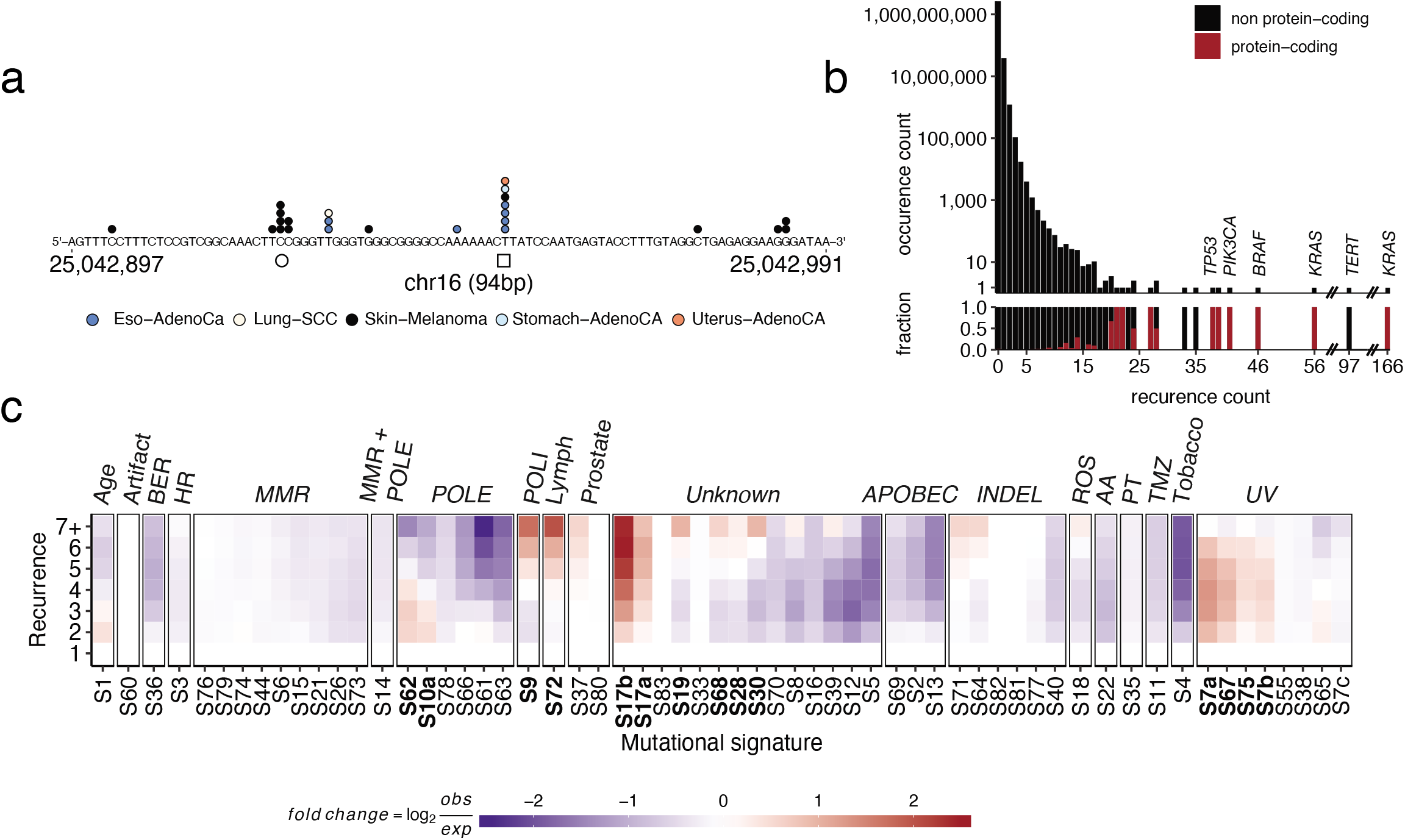
Hotspot overview and identification of enriched localized mutational processes. (a) Pan-cancer recurrent and singleton SNVs in a 94-bp window on chromosome 16. Two cases of 11-mer instances containing hotspots. SNVs are colored by cancer type, (b) Hotspot recurrence counts (x-axis) and frequency in counts (y-axis; top) with the proportion (bottom) of positions in coding (red) or non-protein-coding regions (black), (c) All SNVs (n = 41,318,716) grouped by their pan-cancer recurrence count (1-7+). Heatmap showing the relative contribution of all mutational signatures (x-axis) to mutations of hotspots of increasing recurrence (y-axis). Colors represent log2-fold change in mean contribution relative to singleton SNVs (recurrence 1). Several mutational signatures are enriched (red) in highly recurrent hotspots (recurrence 5, 6, 7+).

Highly recurrent hotspots, where ≥25 genomes share the mutation, are mainly found in protein-coding regions (62% [8 out of 13]; **Fig. 3b**). These include drivers in known cancer genes such as *KRAS, BRAF*, and *TP53* [57] and they are the results of recurrent positive selection [7]. We omit these from our analysis, as they are primarily a result of recurrent selection rather than shared localized mutation processes [5, 7].

We next asked whether any mutational signatures were enriched at hotspots, which would suggest they captured localized mutational processes with strong context preferences. For this, we evaluated the contribution of each mutational signature to the mutations of each hotspot. We then divided the hotspots into recurrence classes, where recurrence class one represents SNVs outside of hotspots, so-called singletons. We found that several mutational signatures of both known and unknown etiologies were enriched among hotspots and that the enrichment often increased with recurrence (**Fig. 3c**). Specifically, we found that the signature 17b signal in highly recurrent (5, 6, 7+) SNVs was 6.4-fold enriched from singletons. We also found hotspot-enriched signatures related to UV (signatures 7a, 67, 75, 7b), POLE (62, 10a), POLI (9), and linked to lymphoma (72) as well as several of unknown etiologies (17b, 17a, 19, 68, 28, 30).

Using the full dataset, we compared mutation rates across nested 11-mer subsets with increasing recurrence (**Fig. 4a**): a set of 11-mers that harbor at least one (1+) singleton (n = 1,796,253 11-mers), a set of 11-mers with mutations in two or more (2+) genomes (n = 351,996 11-mers), and a set of 11-mers mutated in five or more (5+) genomes (n = 3,817 11-mers). The genomic span of these 11-mer sets were 712- (2+; 954 Mb/1.3 Mb) and 3,813-times (5+; 23 Mb/6.2 Kb) higher than the hotspot positions used to define them. The mutation rate of the hotspot set (2+; 10.02 SNV/patient/Mb) was 1.7x increased over the full 11-mer set (1+; 5.96 SNV/patient/Mb), while the highly recurrent hotspots (5+; 25.53 SNV/patient/Mb) set had 4.3x increased mutation rates.

**Figure 4.**
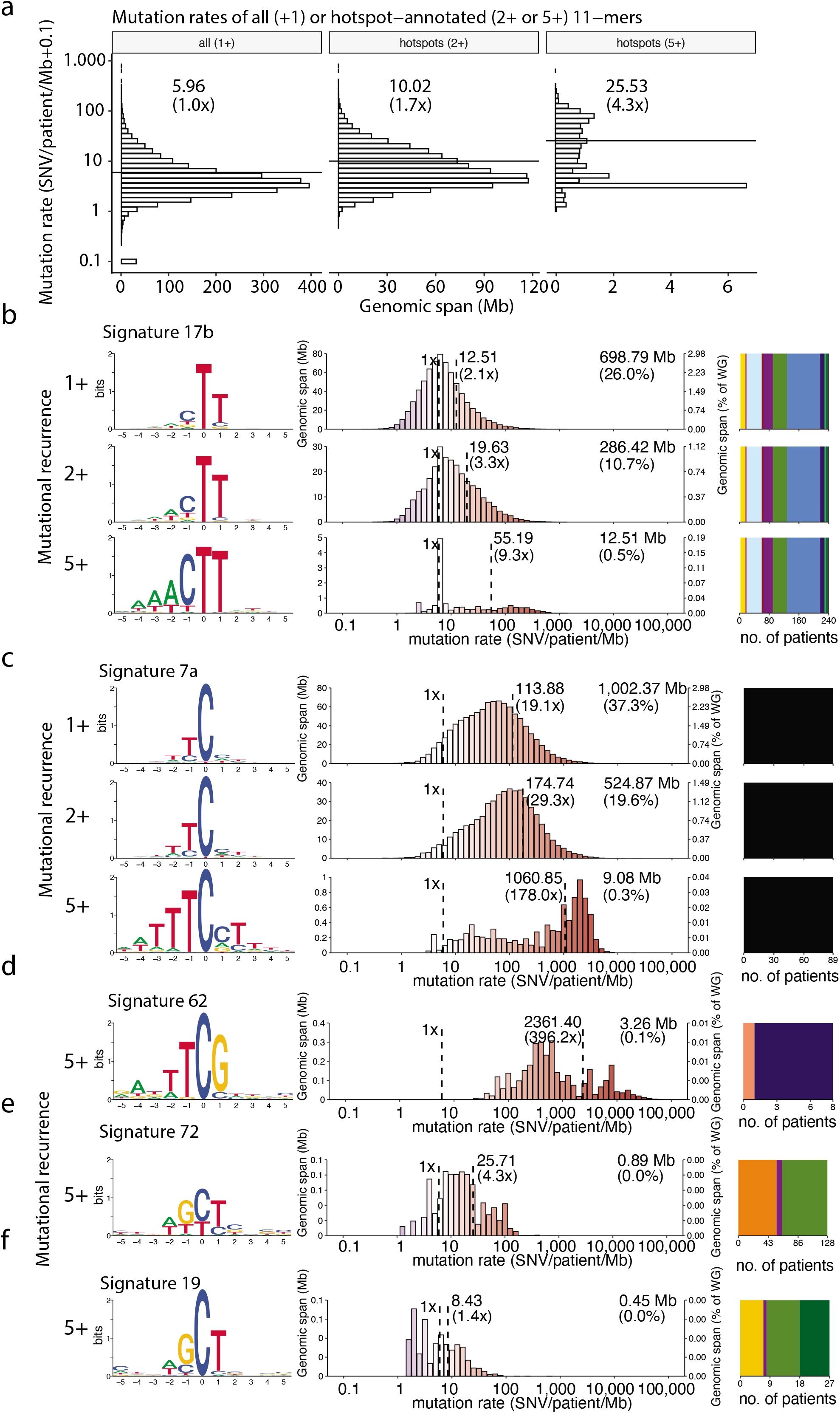
Mutational recurrence capture highly mutated 11-mers. (a) Mutation rates of all mutated 11-mers (1+; 98.8% [2,653 Mb] of the genome) and 11-mers with a hotspot in at least one of its instances for all hotspots (2+; 35.5% [954 Mb] of the genome), and highly recurrent hotspots (5+; 0.9% [23 Mb] of the genome), (b) Signature 17b-assigned 11-mers of all recurrences-levels (1+; top horizontal panels), 11-mers with a hotspot in at least one of its instances (2+; middle horizontal panels), and 11-mers with a highly recurrent hotspot in at least one of its instances (5+; bottom horizontal panels). Logo plots (left) visually highlight contexts different from the background base distribution and thus have the potential to capture sequence dependencies. Histograms (middle) show the distribution of mutation rates of the 11-mers subsets. Stacked bar plots (right) represent the cancer type distribution, colored as in Fig.1a. (c) Plots as in (b) with 11-mers assigned to UV-signature 7a. (d) POLE-signature 62-assigned 11-mers with a highly recurrent hotspot in at least one of its instances (5+). Plots as in (a), (e) Signature 72-assigned 11-mers with a highly recurrent hotspot in at least one of its instances (5+). Plots as in (a), (f) Signature 19-assigned 11-mers with a highly recurrent hotspot in at least one of its instances (5+). Plots as in (a).

When we held out the hotspot mutations used to identify the included 11-mers, the mutation rates were still elevated by 1.6x for the 2+ set and by 4.5x for the 5+ set (**Suppl Fig. 1**), which shows that the high mutation rates of these 11-mers are not simply the result of ascertainment bias and that the higher rates are also driven by singletons. Thus, hotspots enable us to capture highly mutable 11-mer families.

### Characterization of mutational signatures enriched at hotspots

We applied the recurrence-stratification on signature-assigned 11-mer sets. For signature 17b-assigned 11-mers with high recurrence levels (5+), we found a 9.3-fold enrichment in mutation rate and strong enrichment of adenines in the three 5’-positions offset (fourth, third, and second neighbor) from the mutated base (AAAC<u>T</u>T; **Fig. 4b**). When we accounted for the nucleotide composition bias from the mutational signature profile (**Suppl. Fig. 5**; Methods), the 5’-A-tract remained highly enriched (**Suppl. Fig. 4**). A subset (AAC<u>T</u>T) of this motif has also been reported by Stobbe *et al*. (2019) [21], while Alexandrov *et al*. (2020) [22] showed high mutation type probabilities in AC<u>T</u>TA when fitting to pentanucleotide signatures. The wide range of cancer types affected by this signature in the PCAWG dataset includes adenocarcinomas of the digestive system (esophagus, stomach, colorectum, pancreas, and biliary bladder; n=170), breast (n=4), and lung (n=3), as well as B-cell non-Hodgkin lymphoma (BNHL; n=38), bone osteosarcoma (n=13), head and neck squamous cell carcinoma (n=4), hepatocellular carcinoma (n=4), skin melanoma (n=3), and chromophobe renal cell carcinoma (n=1).

We also found that the UV-associated signature 7a was enriched in hotspots (**Fig. 3c**), and the mutation rate of signature 7a-assigned 11-mers with 5+ hotspots was enriched 178-fold compared to the baseline mutation rate (**Fig. 4c**). The nucleotide composition of this 11-mer subset displayed trends towards the T<u>C</u>S (S=C|G) center trinucleotide flanked by additional up- and downstream thymines (TTT<u>C</u>ST). This motif has previously been reported [21, 29, 30]. While the emergence of this motif is driven by highly mutated 11-mers with mutation rates above the mean (164 SNV/patient/Mb), we observed a different nucleotide composition in the lowly mutated contexts (WS<u>Y</u>T; W=A|T, Y=C|T; **Suppl. Fig. 2**).

In genomes from adenocarcinoma of the colorectum and ovary (n=8), 11-mers with 5+ hotspots assigned to the mutational signature 62 of POLE deficiency displays specificity toward the TT<u>C</u>G motif at mutation rates 396-fold higher than the baseline (**Fig. 4d**). From a pentanucleotide signature model, Alexandrov *et al*. (2020) [22] showed that signature 62 has moderate preference towards C>T substitutions in a TT<u>C</u>G context, however they found that C>A substitutions in TT<u>C</u>TT were much more likely for this signature. The TT<u>C</u>G context has also been reported by others [22, 58, 59]. Our findings suggest that POLE-associated signature 62 displays highly localized mutagenesis in TT<u>C</u>G contexts. We also found highly increased mutation rates and strong sequence specificities towards the TTT<u>C</u>TTT hepta-nucleotide motif for POLE-signatures 10a (265-fold) and 61 (184-fold; **Suppl. Fig. 4**). This is an extension of the highly mutable TT<u>C</u>TT motif modeled by the POLE-associated pentanucleotide signatures 10a, 61, 62, 63, and 66 from Alexandrov *et al*. (2020) [22].

For signature 72, which is associated with B-cell lymphomas (BNHL and chronic lymphocytic leukemia), we observed 4.3-fold increased mutation rates in the 5+ set over the baseline (**Fig. 4e**). The nucleotide context showed a strong trend toward the WG<u>C</u>T motif. Though signature 72 has no clear etiology, this motif highly resembles a known hotspot motif (AG<u>C</u>T) of AID activity [60, 61], known to be involved in lymphomagenesis [62].

The AID hotspot motif also emerged from the 5+ set assigned to signature 19, and the mutation rates increased 1.4-fold over the baseline (**Fig. 4f**). Signature 19 is active in BNHL genomes, but no etiology has been proposed for this signature. Though the mutational profile of signature 19 is very different from signature 72 (cosine similarity = 0.24), the similar sequence contexts of these signature-assigned 11-mers with hotspots support a relatedness to AID-mutagenesis.

### Localized mutational processes are operative in distinct genomic elements

To evaluate whether the hotspot-associated mutational processes show preference for specific genomic regions, we examined the mutation rate of signature-assigned 11-mers found within functional genomic elements from ENCODE [63] and compared them to the equivalent subsets of genome-wide 11-mers. We expected the mean mutation rate of 11-mers in each genomic region to be equal to that of the genome-wide subset when genomic regions do not affect mutagenicity. Contrarily, we found that certain genomic regions contain 11-mers with higher mutation rates compared to the corresponding genome-wide subset (**Fig. 5**).

**Figure 5.**
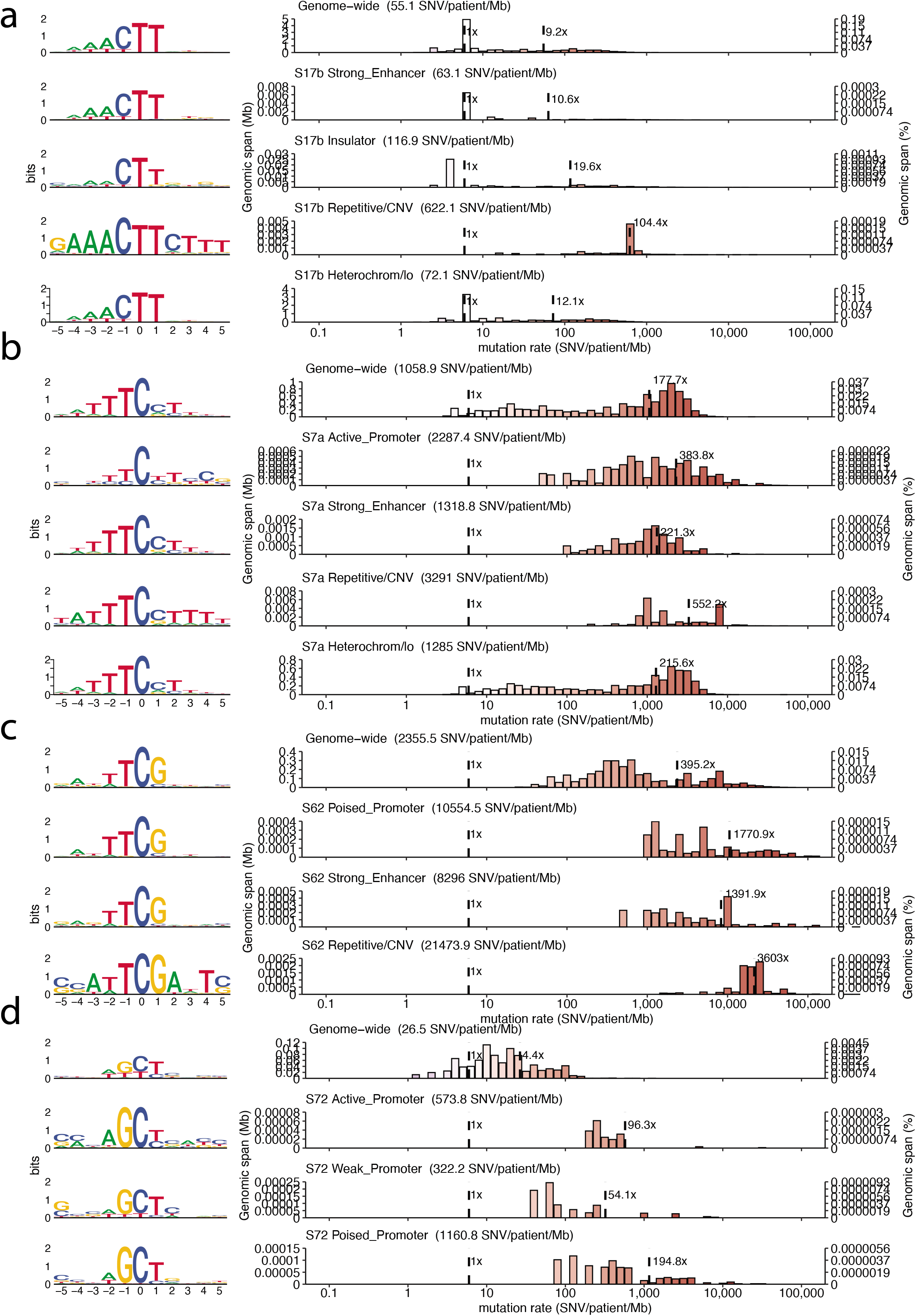
Genomic regions with highly mutable 11-mers. (a) Sequence characteristics (left; logo plot) and mutation rate distribution (right; histogram) of signature 17b-assigned 11-mers of high recurrence (5+) across the whole genome (top) and most highly mutated genetic elements (below), (b) Signature 7a-assigned 11-mers of high recurrence (5+) characterized as in (a), (c) Signature 62-assigned 11-mers of high recurrence (5+) characterized as in (a), (d) Signature 72-assigned 11-mers of high recurrence (5+) characterized as in (a).

For signature 17b, the mutation rate of 11-mers drastically increased in enhancers (10.6-fold), insulators (19.6-fold), heterochromatin (12.1-fold), and repetitive regions (104-fold) (**Fig. 5a**). We found that enhancers and heterochromatin displayed weak 5’-A-tracts, while the repetitive regions were strongly enriched for an extended motif (GAAAC<u>T</u>TCTTT; **Fig. 5a**) beyond what is captured by hotspots (AAAC<u>T</u>T; **Fig. 4b**). Interestingly, the same 11-bp sequence context in repetitive regions also showed high mutation rates for POLE signatures 78 (183.7-fold) and 63 (1,120-fold) (**Suppl. Fig. 4**).

To further evaluate GAAACTTCTTT mutability in repetive elements, we annotated 11-mer instances with repeat-classes from RepeatMasker [64] (Methods; **Suppl. Fig. 3**). We found that this 11-mer is indeed highly mutable (72.8-fold) in repetitive regions pan-cancer. Additionally, we observed that the mutated instances almost exclusively (82.8%; 1,200 out of 1,450) occured in alpha satellite repeats, characteristic of the centromeres.

For the UV signature 7a, 11-mer mutation rates increased in heterochromatin (215-fold), enhancers (222-fold), promoters (384-fold), and repetitive regions (552-fold) (**Fig. 5b**). The 11-mer subsets within insulators, enhancers, heterochromatin and repetitive regions had strong sequence tendencies towards the TTT<u>C</u>STT (S=C|G) motif, consistent with previous reports of T-tracts in UV hotspot motifs [26, 65, 66]. This motif was far less pronounced in promoters, even though they have previously been coupled to increased UV-mutability [27, 29].

The POLE-associated (signature 62) subsets displayed strong sequence preferences for the POLE-motif (TT<u>C</u>G) and dramatically increased mutation rates in promoters (1,771-fold), enhancers (1,392-fold), and repetitive elements (3,603-fold) (**Fig. 5c**). The latter showed not only the highest mutation rate, but also strong sequence preference (ATT<u>C</u>GA) for an adenine flanking each end of the POLE-motif.

Last, we found increased mutation rate for the B-cell lymphoma signature 72 in active (96-fold), weak (54-fold), and poised promoters (195-fold), which were further enriched for the motif (AG<u>C</u>T) seen in genome-wide hotspots (**Fig. 5d**). Similarly, signature 19 with the same hotspot-motif, displayed strong sequence dependency and increased mutation rate in poised promoters (202-fold) (**Suppl. Fig. 4**).

### Several signatures exhibit strongly localized behavior

In combination, we identified sets of positions in specific genomic regions that are targeted by localized mutational processes and subject to much elevated mutation rates (**Fig. 6**). We can decompose the increase in mutation rate into explanatory factors. Together, these factors each define increasingly smaller parts of the genome where the underlying processes are increasingly active. This allows us to identify the sequence characteristics of highly mutable contexts and the relative rate increase they contribute.

**Figure 6.**
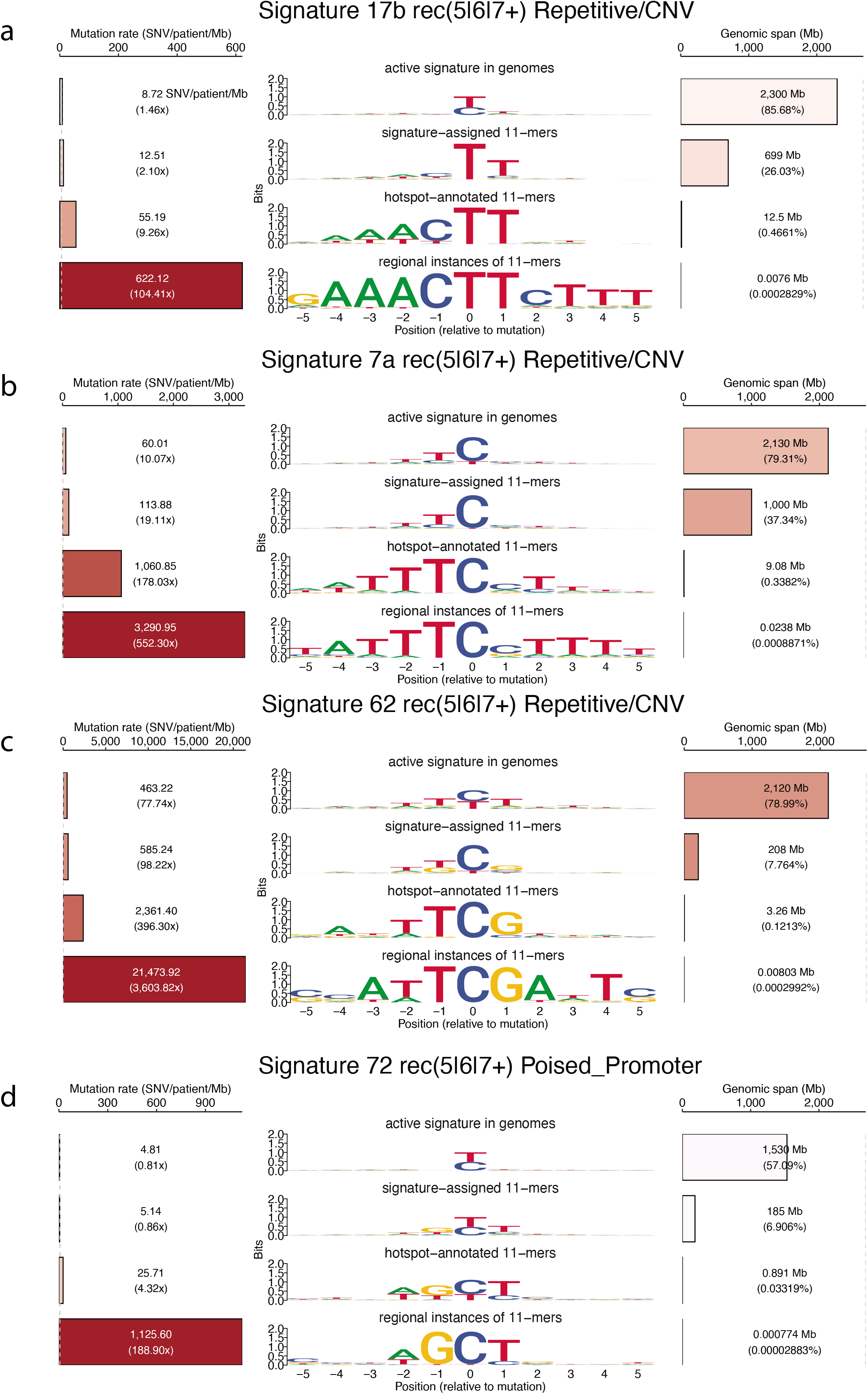
Decomposition of factors increasing the mutation rate. (a) For signature 17b, each analysis step is represented by the mean 11-mer mutation rate (left; barplot), sequence characteristics (middle; logo plot), and genomic span (right; barplot). Mutation rate is quantified as SNV/patient/Mb and the increase from the baseline mutation rate (5.96 SNV/patient/Mb) is stated in parenthesis. For logo plots, letter height is measured in bits (uniform background). The genomic span is given in mega base pair (Mb) with percent of whole genome in parenthesis, (b) For signature 7a, as in (a), (c) For signature 62, as in (a), (d) For signature 72, as in (a).

For instance, for signature 17b (**Fig. 6a**), the exposure-cohort has a modestly increased mutation rate over the baseline (1.5x rate increase; 2,300 Mb genomic span), which is further increased for the large context set where it dominates (1.4x; 699 Mb). Recurrently mutated contexts (4.4x; 12.5 Mb) and repetitive regions (11.3x; 7.6 Kb) further restrict the set of positions to a well-defined 11-bp context (GAAAC<u>T</u>TCTTT) with a dramatically elevated mutation rate (104-fold). This mutational signature has been associated with gastrointestinal cancers and exposure to the genotoxic chemotherapy 5-fluorouracil, though no explanation exists for increased mutability in this highly defined nucleotide sequence [67]. Where available (136 out of 240 patients), the clinical data showed that no patients were exposed to neoadjuvant chemotherapy, thus these tumors are treatment naive and we can rule out 5-fluorouracil as the explanatory process for them.

Samples exposed to the main UV-signature (7a) generally have high mutation rates (10x; 2,130 Mb). When further restricted to contexts where the signature dominates (1.9x; 1,000 Mb), contexts with mutational recurrence (9.3x; 9.1 Mb), and finally repetitive regions (3.1x; 23.8 Kb) the mutation rate increases at scales similar to signature 17b (**Fig. 6b**). Despite their differences in exposed tissues, the processes underlying signatures 7a (UV) and 17b (unknown) both prefer sequence motifs with A/T-tracts 5’ to the mutated nucleotide at similar rates.

Generally, patients exposed to POLE-signature 62 had very high mutation rates (77.7x; 2,120 Mb) with high fractions (median exposure 17.9%) of mutations explained by this signature (**Fig. 6c**). Consequently, signature 62-contexts increased only slightly in mutation rate over the exposed cohort (1.3x; 206 Mb). Extending on signature contribution, both high mutational recurrence (4.0x; 3.3 Mb) and location in repetitive regions (9.1x; 8.0 Kb) contributed large mutation rate increases. Compared to signature 62, mutational recurrence contributed slightly less to the mutation rate in POLE-associated signatures 10a (2.0x; 3 Mb) and 61 (1.6x; 15 Mb). However, for the highly mutables contexts, POLE-signatures 10a and 61 showed preference for a different core motif (TT<u>C</u>T) than for signature 62 (TT<u>C</u>G) (**Suppl. Fig. 4**). This may reflect that POLE deficiency can lead to distinct mechanistic processes.

While the mutational signature 72 by itself did not result in dramatic mutation rate changes, mutational recurrence provided an increased rate (5.0x; 0.89 Mb) similar to the effect seen in the above examples (4-9x). Signature 72 and 19 of unknown etiologies shared the preference for a common motif (AG<u>C</u>T) known as the AID-hotspot motif [60, 61].

In the four cases above, hotspots contributed with a 4-9x increased mutation rate over mutational signatures, which is consistent with our signature-agnostic hotspot-characterization (4.3x; **Fig. 4a**).

## Discussion

In this study, we exploited mutational hotspots to define subsets of the genome that are targeted by localized mutational processes and systematically catalog their mutation rates and sequence preferences. We found that mutation rates increase by 4-400 fold compared to the average pan-cancer mutation rate (baseline) in sequence contexts subject to localized mutational processes associated with UV (signature 7a), POLE (signature 62), lymphomas (signature 72), and an unknown etiology (signature 17b). This is 5-18 times higher than what can be explained by cancer type and mutational signatures alone. Additionally, we found that mutation rates are further elevated (104-3,604 fold) in distinct sequence motifs within genomic regions related to repetitive DNA (signatures 17b, 7a, 62) and promoters (signature 72). We provide a comprehensive catalog of localized mutational processes, their sequence motifs, and their observed mutation rates (**Suppl. Fig. 4**).

Consistent with literature, we found that UV-associated mutagenesis (signature 7a) targets TTT<u>C</u>ST-sequences (S=C|G), which are highly mutated across multiple genomic regions [21, 26, 30]. However, the highly mutated contexts are more ambiguous in promoters, and thus we did not observe a clear motif for these regions. Interestingly, melanoma genomes frequently harbor hotspot mutations in promoter elements explained by ETS-mediated sensitization of DNA to UV-induced cyclobutane pyrimidine dimer formation [27, 28, 68, 69]. The binding of DNA by ETS-transcription factors is estimated to contribute a 16-170-fold elevated mutation rate at ETS-binding sites (CTT<u>CC</u>GG and <u>YY</u>TTCC) [28, 69]. We did not observe this ETS-motif in our analyses. However, for UV-assigned 11-mers with high recurrence, we found a bimodal distribution of mutation rates associated with different sequence preferences (TTT<u>C</u>ST [high] and WS<u>Y</u>T [low]), thus potentially capturing multiple mechanisms by which UV may induce mutations. This shows that our k-mer-centric and rate-based analysis approach can aid in the generation of mechanistic hypotheses for mutational processes. Similar approaches will gain increased power in future large whole-genome cancer datasets.

We observed that two signatures of unknown etiology (signatures 19 and 72) are associated with a hotspot motif (WG<u>C</u>T), which highly resembles the known AID hotspot motif (AG<u>C</u>T) [60, 61]. Additionally, these processes have increased mutability in promoters, which is in line with reported AID off-target effects [70]. Thus, the potential of capturing AID-mutagenesis through signatures 19 and 72 may be further explored.

We found that the rate of signature 17b-mutations is elevated (9-fold) in a genome-wide hotspot motif (AAAC<u>T</u>T) (**Fig. 4b**), which adds more context to the previously identified signature 17-motifs (AC<u>T</u>TA and AAC<u>T</u>T) [21, 22, 71].

Consistent with signature 17 mutations being enriched in cohesin/CTCF-binding sites [72–74], we found a 20-fold mutation rate increase in certain contexts within insulator elements (**Fig. 5a**). However, in these regions, we did not observe the signature 17b-characteristic 5’-A-tract before the C<u>T</u>T core nucleotides. Thus, the mutational mechanism acting in these elements may be distinct from those causing AAAC<u>T</u>T-hotspot mutations in the rest of the genome.

Unexpectedly, we also found a highly enriched 11-mer (GAAAC<u>T</u>TCTTT) in the alpha satellite repeats of centromeric regions, which was associated with both signature 17b and the POLE signatures (63 and 78). This 11-mer contains the reported 5’-A-tract, however it also contains some intrinsic repeat structure that may be broken down into triplicates of the repeat unit, S(W)_2-3_ (S=C|G; W=A|T). Such repeats may adopt secondary DNA structures that facilitate mutagenesis by certain processes, such as APOBECs targeting single-stranded DNA in stem-loops [36, 38, 40] or MMR deficiency leading to increased mutability of AT-rich short inverted repeats [39]. As alpha satellite repeats are replicated in the late S-phase [75], the mutational processes shaping this part of the genome are likely linked to late replication. Mutagenesis from POLE deficiency and the signature 17 process are both associated with late replication [36, 52]. Taken together, this is consistent with GAAAC<u>T</u>TCTTT being associated with these processes in our analyses.

Just like the other motifs subject to tissue-specific localized mutational processes, the AAAC<u>T</u>T motif possesses properties that either increase susceptibility to DNA damage, avoidance of repair, or both. Replication-timing and strand-asymmetry profiles of signature 17-mutations have been shown to be similar to those found for signatures of tobacco and UV exposure. Thus, they may share the property of being linked to environmental DNA-damage mechanisms [52]. Specifically, oxidative damage to the dGTP pool has been proposed as a possible explanation for signature 17-mutations, resulting from exposure to gastric acid in gastrointestinal tumors or exposure to the genotoxic chemotherapeutic 5-fluorouracil in treated tumors [19, 52, 71, 76]. However, these hypotheses do not explain the characteristic motif of signature 17-mutagenesis and the mechanisms involved remain largely unexplained [67].

The signature 17-mutational process has been shown to correlate with the helical periodicity of DNA wound around the nucleosome core [77]. The highest mutation rates are found in the nucleosome-facing minor grooves, likely explained by hindered base excision repair in these sites [77]. While the rigid structure of long A-tracts may constrain DNA winding around the nucleosome [78], short A-tracts likely affect nucleosomal DNA flexibility and thus direct their positioning within the nucleosome with respect to the dyad [79, 80]. Such intra-nucleosomal forces may in turn hinder DNA repair at nucleosome-facing minor groove C<u>T</u>T lesions, thus in part explaining the A-tract motif associated with these mutations. At least, it is possible that lesions in proximity of A-tracts are repaired at different rates than the rest of the genome [81].

In agreement with existing literature [21, 22, 58, 59], we found POLE-mutagenesis to be associated with two highly mutated motifs (TT<u>C</u>G and TTT<u>C</u>TTT) and that their mutation rates dramatically increased over the baseline (184-396-fold). Mutations localized to the TT<u>C</u>G motif seem to be more pronounced for signature 62 than any other POLE signature, though this signature also encompasses T<u>C</u>T mutations. Fang *et al*. (2020) [59] suggest that mutations acquired in distinct domains of the POLE gene may give rise to distinct mutational patterns depending on the mutant-POLE DNA-affinity. Thus, it is possible that there exists even more examples of single mutagenic mechanisms generating different mutation types dependent on their specific loss- or gain-of-function mutants.

## Conclusion

Our findings provide higher resolution of the sequences targeted by localized mutational processes and contribute mutation rate estimates of these. Our comprehensive catalog (**Suppl. Fig. 4**) of mutational processes may aid the construction of more accurate models of the mutational processes in cancer, which capture the mutation rate variation. Such models are important for accurate statistical driver identification among the landscape of passenger hotspot mutations caused by localized processes [82]. In addition, the models may also contribute to deeper understanding of cancer risk, somatic evolution, cancer development, and tumor biology.

The mutational patterns of localized processes active across cancers may serve as future biomarkers for detection of such processes and their associated etiologies in cancer samples. In samples with weak mutation signals, catalogs of localized mutational processes may power detection of active processes through targeted sequencing of their possible genomic targets. For cancer-associated mutational processes, this may translate to new opportunities for liquid biopsies to enable early cancer detection and surveillance of cancer evolution in the patient.

## Methods

### Whole cancer genome data set

The analysis was based on the full set of SNVs calls of 2,583 cancer genomes calls generated by the The Pan-Cancer Analysis of Whole-Genomes (PCAWG) consortium [6]. The GRCh37/hg19 reference genome was used throughout.We focused on SNVs in the non-protein-coding part of the autosomal chromosomes. We excluded protein-coding regions to reduce potential signals of positive selection.The sex chromosomes were excluded as they include a higher rate of false SNVs calls [6].

### Counting k-mer occurrences

First, we counted the number k-mer instances in chromosome 1-22 using the oligonucleotideFrequency function from the Biostrings (version 2.50.2) package in R (version 3.5.1). We obtained the chromosome sequences through the R package BSgenome.Hsapiens.UCSC.hg19 (version 1.4.0). Second, we summed the counts of identical k-mers across the chromosomes. Third, to achieve strand symmetry, we collapsed reverse complementary pairs of k-mers and represented them by the sequence with a center pyrimidine (C or T) together with the total pair sum. For example, for k=11, the AAAGAAGTTTC (*n_purine_* = 5,250) and GAAACTTCTTT (*n_pyrimidine_* = 5,495) pair was represented by GAAACTTCTTT (*n_total_* = 10,745).

### Mutational signature annotation

#### Genome-wide mutational signature annotation

Signature posterior probabilities for the 96 different trinucleotide mutation types in each genome were calculated with SignatureAnalyzer and provided by the PCAWG consortium [22]. We downloaded 60 mutational signature annotations of all 2,583 whitelisted PCAWG genomes (www.synapse.org/#!Synapse:syn11761189.6), which describe the exposure to signature X in genome Y We classified a genome as exposed to a given mutational signature, when the signature load was equal to or above 5% of the genome’s mutation burden.

#### SNV-level mutational signature annotation

We assigned the signature posterior probabilities to each mutation, which were annotated with the most likely signature as in [7].

#### 11-mer-wide mutational signature annotation

To further focus our analyses on sequence contexts explained by the mutational signature used to define the signature-exposure cohort, we assign 11-mers to the signature that primarily explains all 11-mer instances with SNVs.

The principle of signature assignment of 11-mers relies on three steps:

1. We calculated the mean posterior signature probabilities across identical 11-mers within an exposure cohort. We averaged posterior signature probabilities of SNVs in hotspots for each hotspot position to yield a position-wise mean, which we then used to calculate the mean across instances of an 11-mer family. This captures the average predicted probability that a given mutational signature generated the mutations.
2. We annotated each 11-mer with the signature that had the highest mean posterior probability and referred to this signature as most likely to explain this set of SNVs.
3. We identified the 11-mer families annotated with the signature in question for the given signature-exposed subset.

### Definition of hotspots and recurrently mutated 11-mers

We used SNV recurrence to identify 11-mers with high expected mutability. A recurrence count for hotspots was defined as the number of pan-cancer genomes with a shared position-specific SNV. We annotated 11-mers with the highest recurrence count observed across its instances. This annotation was used to further subset 11-mers into two groups: (1) 11-mers where at least one instance had a hotspot, i.e. recurrence of two or more, (2+ k-mer set) (2) 11-mers where at least one instance had a hotspot of recurrence five or more (5+ k-mer set).

### Genomic regions

We annotated the mutated 11-mer instances with which genomic region they occurred in and stratified 11-mers according to 15 different regions defined by ENCODE [83, 84]. Further characterization of repeat elements was performed using RepeatMasker (http://www.repeatmasker.org/) [64].

### Mutation rate calculation

For each 11-mer its mutation rate (SNV / patient / Mb) was calculated as follows

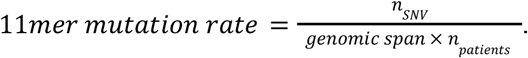

For a set of 11-mers, the mean mutation rate was calculated as follows

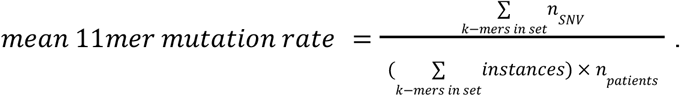

### Sequence context

Sequence information in logo plots was calculated as the Kullback-Leibler divergence between the observed and expected frequency of nucleotides at each position. The expected distribution was derived as the genome-wide autosomal distribution of nucleotides, i.e. A = 29.5%, T = 29.5%, C = 20.5%, and G = 20.5%.

The surprise of observing a nucleotide, *a*, at a given position, *i*, is estimated as the Kullback-Leibler divergence:

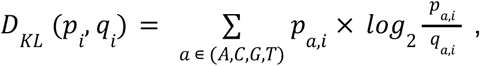

where *p_a,i_* is the observed frequency and *q_a,i_* is the expected frequency of nucleotide *a* in position *i*. The divergence is visualized using a logo plot with letter *height_a,i_* proportional to letter frequency, *p_a,i_*, and divergence, *D_KL_*(*p_i_, q_i_*), in that position:

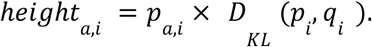

#### Null model of 11-mer nucleotide composition mutational signatures

A null model of 11-mer nucleotide distribution was derived by weighting the genomic 11-mer distribution by the mutation context probability of each signature.

We thereby derived an expected nucleotide distribution in 11-mers under a null model for each signature (**Suppl. Fig. 5**). To achieve this, we more specifically (1) created position frequency matrices for each 11-mer family, (2) collapsed and summed base counts across families sharing trinucleotide context, and (3) weighted each trinucleotide group by the mutation type probability from a given signature.

## Author contributions

Gustav Alexander Poulsgaard (GAP), Simon Grund Sørensen (SGS), Randi Istrup Juul (RIJ), Morten Muhlig Nielsen (MMN), and Jakob Skou Pedersen (JSP).

JSP conceived the project. GAP performed data analysis with contributions from SGS, RIJ, and MMN. GAP drafted the manuscript and all figures. JSP supervised the project. All authors discussed the results and contributed to the final version of this manuscript.

## Acknowledgement

We thank the Independent Research Fund Denmark | Medical Sciences (8021-00419B), Aarhus University, Aarhus University Research Foundation (AUFF-E-2020-6-14), the Danish Cancer Society (R307-A17932), and the Health Research Foundation of Central Denmark Region (R56-A2972-B1845) for funding the project.

We thank the Pan-Cancer Analysis of Whole Genomes (PCAWG) project under the International Cancer Genome Consortium (ICGC) for data access, including variant calls and mutational signatures.

All computing was performed at the GenomeDK high-performance computing (HPC) facility. We thank GenomeDK and Aarhus University for providing HPC resources and support.

## Ethics declarations

Not applicable, as all data comes from the published and fully consented PCAWG study [6].

## Competing interests

The authors declare no competing interests.

## Notes

### Competing Interest Statement

The authors have declared no competing interest.

