## Supplementary Figures 1, 2, 3, 5, and Supplementary Table 1 for "Sequence dependencies and mutation rates of localized mutational processes in cancer"

Mutation rate correction of all (+1) or hotspot−annotated (2+ or 5+) 11−mers

Quantification of ascertainment bias in mutation rate of hotspot−selected 11−mers

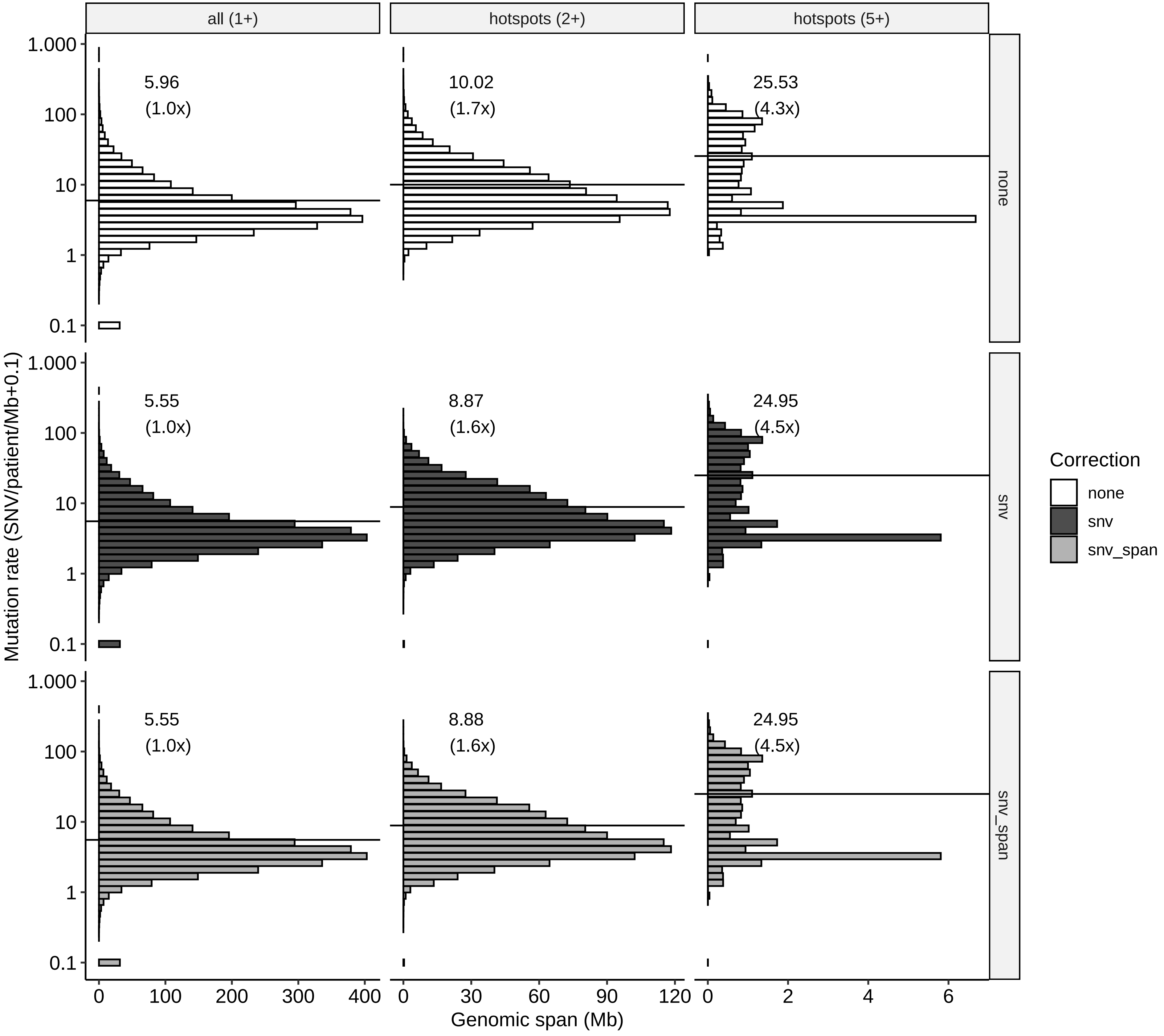


Mutation rate correction method:
none (mut.rate = all SNVs / all 11−mer instances)
conservative (mut.rate = less recurrent SNVs / all 11−mer instances)
fair (mut.rate = less recurrent SNVs / less recurrent 11−mer instances)

**Supplementary Figure 1. Mutation rate estimates of 11-mers with hotspots.**

(Upper horizontal panel) 11-mer mutation rates from all SNVs (1+; left), 11-mers with hotspots (2+; middle), and 11-mers with highly recurrent hotspots (5+; right). (Middle horizontal panel) Mutation rate correction substracting hotspot-SNVs from the mutation rate, for hotspots with equal to or higher recurrence levels than used for selection of the 11-mers. (Lower horizontal panel) Mutation rate correction substracting hotspot-SNVs and genomic span of hotspots with equal to or higher recurrence levels than used for selection of the 11-mers.

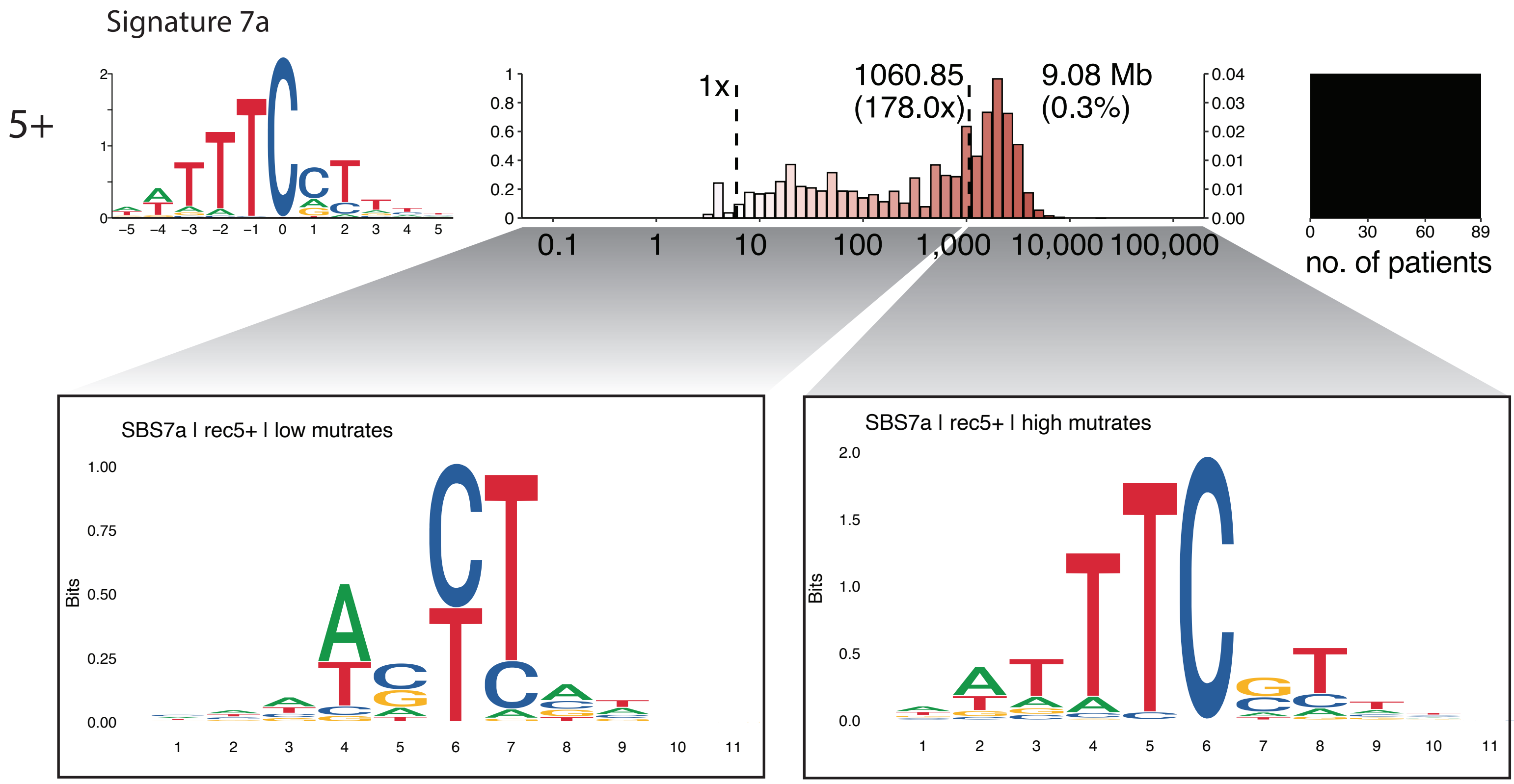


**Supplementary Figure 2. Sequence characteristics across mutation rate distribution.**
From Figure 4c: Signature 7a-assigned 11-mers with hotspots (5+) display a bimodal mutation rate distribution. The sequence characteristics of lowly and highly mutated 11-mers are shown as logo plots.

Suppl. Figure 3

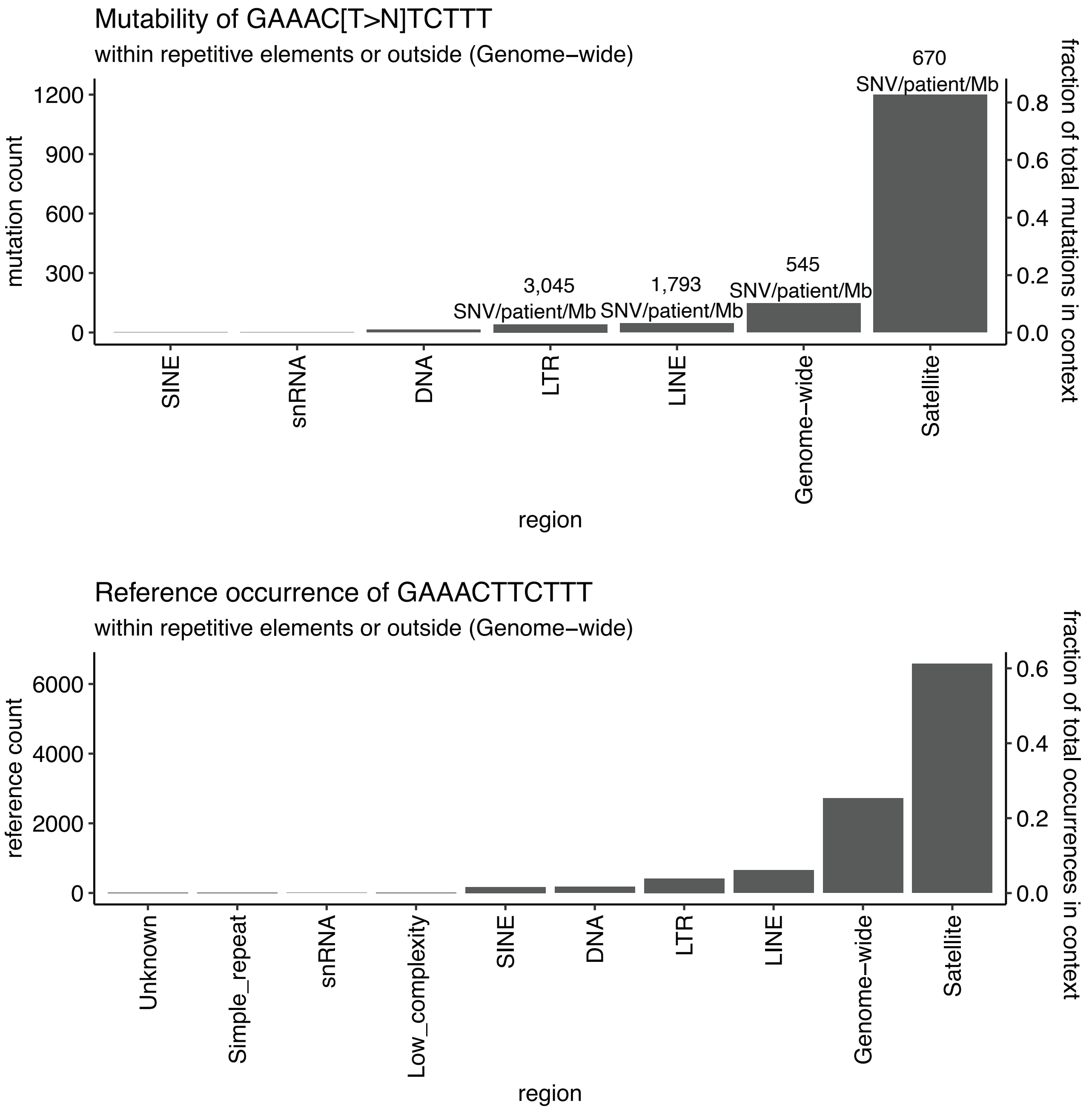


**Supplementary Figure 3. GAAACTTCTTT sequence mutability and occurrence in repetitive regions.**

(Upper plot) The number of mutations (y-axis left) in the GAAACTTCTTT contexts across repetitive elements (RepeatMasker) or outside repetitive elements (Genome-wide). (Lower plot) The occurrence (y-axis left) of GAAACTTCTTT instances across repetitive elements (RepeatMasker) or outside repetitive elements (Genome-wide).

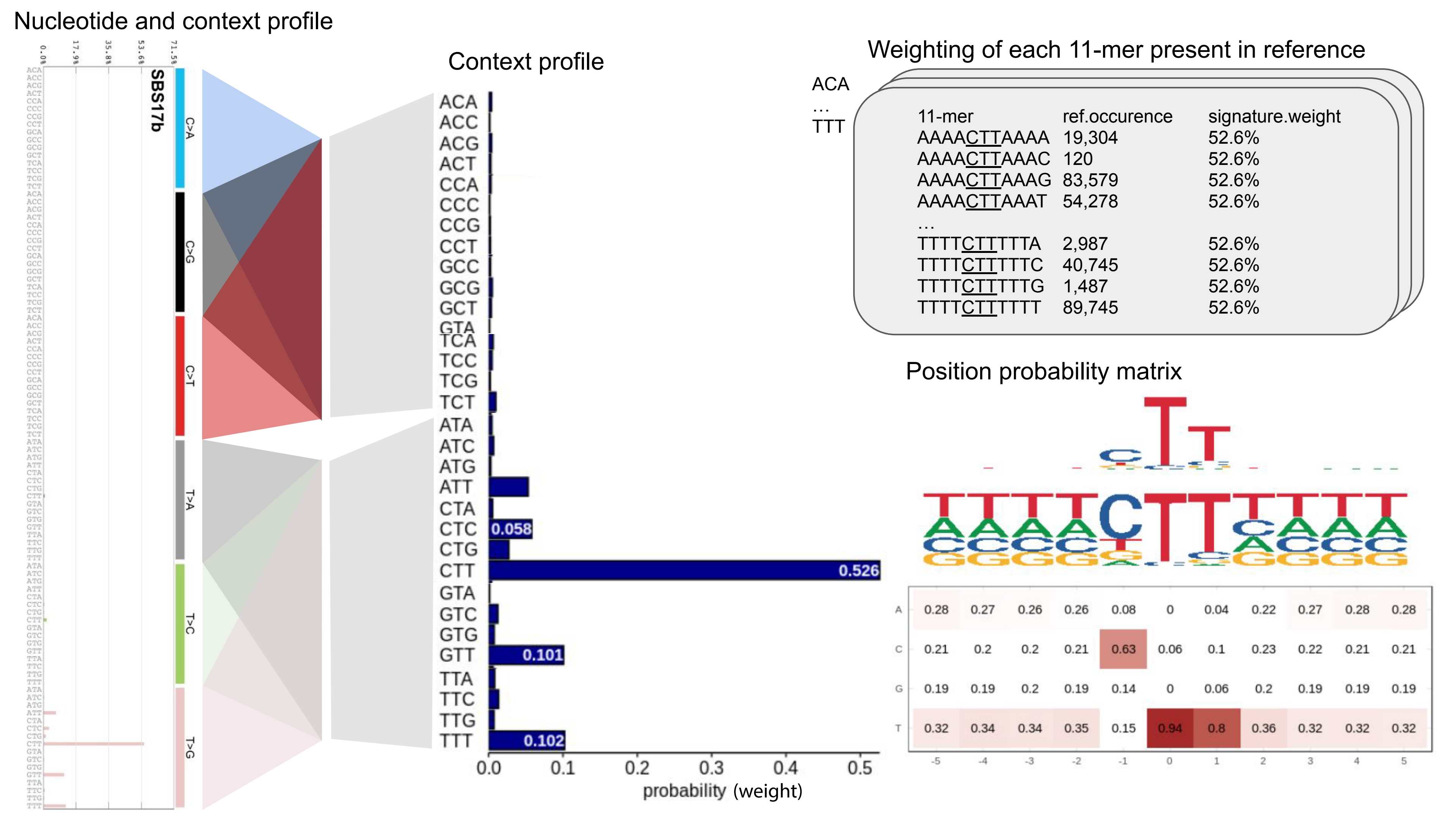

Nucleotide and context profile
SBS17b
C>A
C>G
C>T
T>A
T>C
T>G
Context profile
ACA
ACC
ACG
ACT
CCA
CCC
CCG
CCT
GCC
GCG
GCT
GTA
TCA
TCC
TCG
TCT
ATA
ATC
ATG
ATT
CTA
CTC
CTG
CTT
GTA
GTC
GTG
GTT
TTA
TTC
TTG
TTT
0.058
0.526
0.101
0.102
0.0
0.1
0.2
0.3
0.4
0.5
probability (weight)
Weighting of each 11-mer present in reference
ACA
…
TTT
11-mer ref.occurence signature.weight
AAAACTTAAAA 19,304 52.6%
AAAACTTAAAC 120 52.6%
AAAACTTAAAG 83,579 52.6%
AAAACTTAAAT 54,278 52.6%
…
TTTTCTTTTTA 2,987 52.6%
TTTTCTTTTTC 40,745 52.6%
TTTTCTTTTTG 1,487 52.6%
TTTTCTTTTTT 89,745 52.6%
Position probability matrix
A 0.28 0.27 0.26 0.26 0.08 0 0.04 0.22 0.27 0.28 0.28
C 0.21 0.2 0.2 0.21 0.63 0.06 0.1 0.23 0.22 0.21 0.21
G 0.19 0.19 0.2 0.19 0.14 0 0.06 0.2 0.19 0.19 0.19
T 0.32 0.34 0.34 0.35 0.15 0.94 0.8 0.36 0.32 0.32 0.32
-5 -4 -3 -2 -1 0 1 2 3 4 5

**Supplementary Figure 5. Null model of 11-mer sequence composition.**
**(**Left panel) Mutational signature profile of 96-trinucleotide mutation types. (Middle panel) Collapsed 96 mutation type contexts to 32 trinucleotide contexts containing the sum of probabilities for each mutation type in identical trinucleotide contexts. (Right top panel) The genomic occurrence of each 11-mer is normalized by the trinucleotide context probability and (Right lower panel) the position frequency matrix containing base distribution per position in the collective set of genome-wide 11-mers is computed.

### Supplementary Table 1

| | kmer space | | | | instances per family | | | | | |
|---|---|---|---|---|---|---|---|---|---|---|
| **k** | **expected** | **observed** | **nullomers** | **snvs per kmer (expected)** | **min** | **Q1** | **median** | **mean** | **Q3** | **max** |
| **1** | 2 | 2 | 0 | 20.659.025,00 | 1.100.439.365 | 1.221.362.933,75 | 1.342.286.502,50 | 1.342.286.502,50 | 1.463.210.071,25 | 1.584.133.640 |
| **3** | 32 | 32 | 0 | 1.291.189,06 | 11.923.725 | 67.881.588,50 | 84.697.455,00 | 83.892.887,72 | 105.985.401,25 | 205.859.180 |
| **5** | 512 | 512 | 0 | 80.699,32 | 147.012 | 2.952.289,00 | 5.344.526,50 | 5.243.304,34 | 6.967.661,00 | 36.208.737 |
| **7** | 8.192 | 8.192 | 0 | 5.043,71 | 3.695 | 63.206,25 | 292.403,50 | 327.706,45 | 457.130,00 | 11.824.057 |
| **9** | 131.072 | 131.072 | 0 | 315,23 | 54 | 2.965,75 | 14.723,00 | 20.481,65 | 28.077,25 | 6.870.011 |
| **11** | 2.097.152 | 2.097.090 | 62 | 19,70 | 0 | 146,00 | 608,00 | 1.280,10 | 1.637,00 | 4.674.610 |
| **13** | 33.554.432 | 32.307.152 | 1.247.280 | 1,28 | 0 | 7,00 | 23,00 | 80,01 | 93 | 3.279.735 |
| **15** | 536.870.912,00 | - | - | 0,08 | - | - | - | 5,00 | - | - |

**Supplementary Table 1. Statistics of different k-mer lengths.**
