## Supplementary figures and images for "Sequence dependencies and mutation rates of localized mutational processes in cancer"

### Supplementary Figure 4

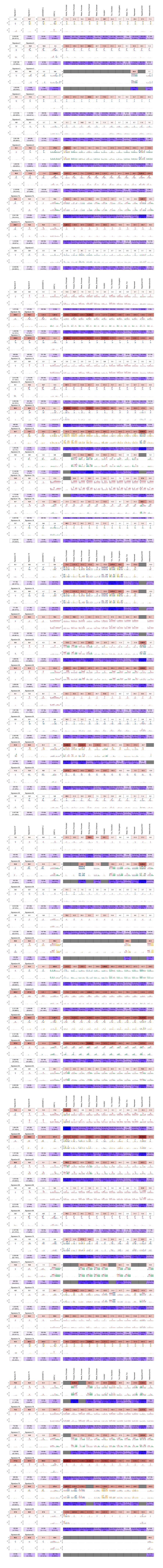
